# Torsional turning motion of chromosomes as an accelerating force to align homolog ous chromosomes during meiosis

**DOI:** 10.1101/459271

**Authors:** Kazutaka Takao, Kazunori Takamiya, Da-Qiao Ding, Tokuko Haraguchi, Yasushi Hiraoka, Hiraku Nishimori, Akinori Awazu

**Affiliations:** Department of Mathematical and Life Sciences, Hiroshima University, 1-3-1 Kagamiyama, Higashi-Hiroshima 739-8526, Japan; Research Center for Mathematics on Chromatin Live Dynamics, Hiroshima University, 1-3-1 Kagamiyama, Higashi-Hiroshima 739-8526, Japan; Advanced ICT Research Institute Kobe, National Institute of Information and Communications Technology, 588-2 Iwaoka, Iwaoka-cho, Nishi-ku, Kobe 651-2492, Japan; Graduate School of Frontier Biosciences, Osaka University, 1-3 Yamadaoka, Suita 565-0871, Japan

## Abstract

Homologous sets of parental chromosomes must pair during meiosis to produce recombined sets of chromosomes for their progeny. This is accompanied by nuclear oscillatory movements. This study aimed to elucidate the significance of these movements with a model, wherein external force was applied to the oscillating nucleus and via hydrodynamic interactions within the nucleus. Simulations revealed that a major force for aligning homologous chromosomes is length-dependent sorting during chromosomal torsional turning, which occur when the nucleus reverses the direction of its movement.

---

Meiosis is an important process for eukaryotic organisms to generate inheritable haploid gametes from a diploid cell. During meiosis, homologous (paternal and maternal) chromosomes recombine; these recombined chromosomes are inherited by their progeny. For recombination of homologous chromosomes, side-by-side alignment of homologous loci along the chromosome is necessary [1]. During this alignment of homologous chromosomes, telomeres form a cluster beneath the nuclear envelope in various eukaryotic organisms [2].

Unicellular fission yeast *Schizosaccharomyces pombe* is a useful model system for studying the alignment of homologous chromosomes during meiosis [3, 4]. A meiotic cell of *S. pombe* contains 3 pairs of homologous chromosomes (chromosomes 1, 2, and 3 of 5.6, 4.6, and 3.5 Mb, respectively) [5]. During meiotic prophase, the nucleus elongates, oscillates between the cell poles, and shows drastic deformation, so-called “horsetail movement” [6], during which telomeres remain clustered at the spindle pole body (SPB) at the leading edge of the nuclear movement [6]. Telomere clustering and nuclear movements reportedly promote the alignment of homologous chromosomes [7,8,9].

To elucidate mechanisms underlying the alignment of homologous chromosomes, computer simulation strongly facilitates the examination and prediction of biological events under conditions that are never accomplished empirically. Various theoretical models of the pairing of homologous loci during meiosis have been proposed on the basis of telomere clustering [10, 11, 12, 13]. However, the role of the nuclear envelope, which confines the chromosomes, remains unknown in these models.

Studies are required to consider the role of the nuclear envelope in models of the meiotic nucleus of *S. pombe*, since its nucleus migrates and undergoes striking deformation during meiosis. During the horsetail movements, the nuclear envelope receives drag forces from the cytoplasm and from the nucleoplasm, while each chromosome receives drag forces only from the nucleoplasm. Thus, models of the meiotic nucleus of *S. pombe* would help describe the motion of the deformable nuclear envelope, direct collisions between the nuclear envelope and chromosomes, effects of the cytoplasm on the nuclear envelope, and effects of the nucleoplasm on chromosomes and the nuclear envelope, in addition to telomere clustering at the SPB. This study aimed to propose the first model displaying the aforementioned nuclear membrane dynamics and investigate the role the horsetail nuclear movements in the alignment of homologous chromosomes via simulations of the presented model..

## Model construction

A model of chromosomes confined within the nuclear envelope of *S. pombe* cells (Fig. 1(a)) was developed using the following methods. We described each chromosome as a chain and the nuclear envelope as a one-layered shell—both comprising particles with an excluded volume (Fig. 1(b)). Each chromosome was assumed to consist of particles with uniform radii since they were expected as almost uniformly condensed chains during meiosis. A meiotic nucleus of *S. pombe* contains 3 homologous pairs of chromosomes; thus, the model contains 6 chains in total confined in the shell. To account for the effects of the SPB on chromosomal movement, we constructed a virtual sphere around the SPB (“the SPB sphere”), penetrating a limited area of the shell (nuclear envelope) extending toward the space inside the shell (Fig. 1(b)). To simulate telomere movement, which are clustered near the SPB during the horsetail movement, we assumed that ends of the chromosome chains move along the surface of the SPB sphere inside the shell.

**FIG. 1.**
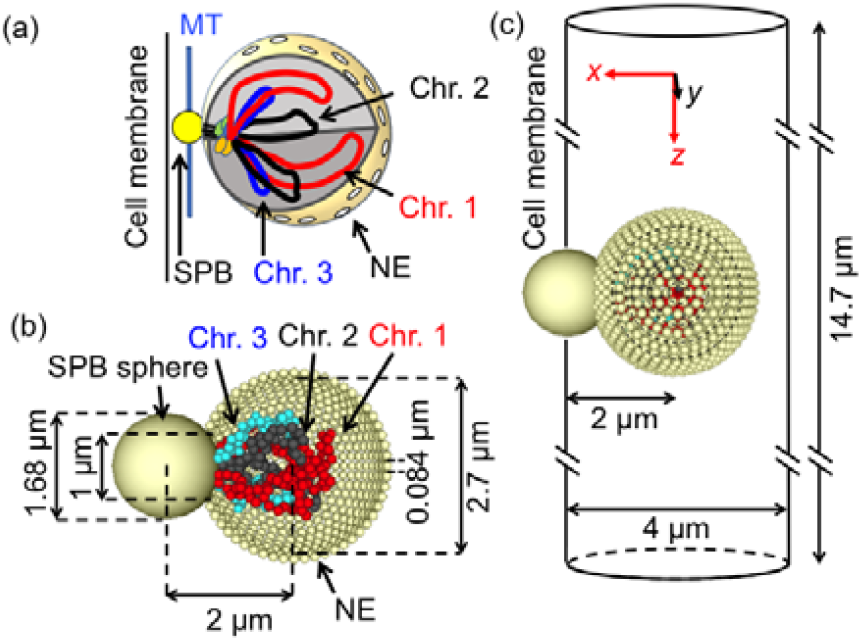
(a) Illustration of the meiotic nucleus of *Schizosacchromyces pombe* containing 3 homologous pairs of chromosomes (Chr, 1, 2, and 3), the nuclear envelope (NE), the spindle pole body (SPB), and microtubules (MT). (b) Illustration of the model composed of particle populations. (c) Illustration of the model of an intracellular region described as a cylindrical space, and the initial structure of the nuclear envelope constructed by combining particle rings (along dashed circles) along the vertical y-axis with various radii.

To simulate the horsetail movements, we assumed that the SPB sphere oscillates periodically in the cell. Here, the drag force from the cytoplasm is exerted exclusively on the nuclear envelope. Since the cytoplasm can be considered a static intracellular fluid, we assumed that this drag force is proportional to the velocity of the nuclear envelope. However, the drag forces from the nucleoplasm to the nuclear envelope and chromosomes depend on the intranuclear velocity profile of the nucleoplasm, which is primarily determined on the basis of velocity profiles of chromosomes and the nuclear envelope. Thus, drag forces from the nucleoplasm could be described by the hydrodynamic interaction among particles in the chains and the shell. Furthermore, we assumed the volume of the nucleus tend to be sustained since the in and out flow of nucleoplasm through the nuclear envelope were considered to be little during meiosis.

These methods allowed for the generation of an appropriate model involving the effects indispensable for simulating the *S. pombe* meiotic nucleus.

## Model implementation

We assumed that chromosomes 1, 2, and 3 consist of *N_n_* particles (n = 1, 2, or 3) and the nuclear envelope consists of *M* particles with radii = *r*. The motion of the *i*-th particle is given as

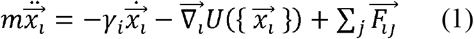

where 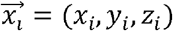 and *m* are the position and mass of the *i*-th particle, respectively. *γ_i_* indicates the coefficient of a drag force from the cytoplasm where *γ_i_* = 6*πrη* if the *i*-th particle is on the nuclear envelope and *γ_i_* = 0 if the *i*-th particle is on the chromosome. Here, *η* was assumed as the viscosity of cytoplasm.

In the second term of equation (1), the potential of the system 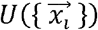 is given as follows:

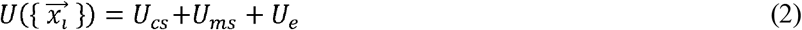

where *U_cs_* is the contribution from the chain, *U_ms_* is the contribution from the nuclear envelope, and *U_e_ is* the potential of the excluded volume effects.

The potential contribution from the chromosome chain *U_cs_* is given as follows:

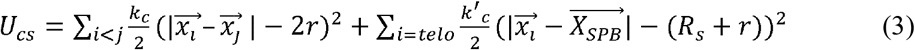

where the *i*-th particle is a part of chromosomes, *r* indicates the radii of particles, and 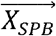 and *R_S_* indicate the position of the center and the radius of the SPB sphere. We assumed each chromosome is a chain of particles connected in a line. Σ in the 1st term indicates the sum for the *i*-th and *j*-th particles (*i* < *j*) that are adjacent and bonded along the chromosome, and Σ in the 2nd term indicates the sum for the *i*-th particle at the ends of chromosomes corresponding to the telomeres.

The potential contribution from the nuclear envelope *U_ms_* is given as follows:

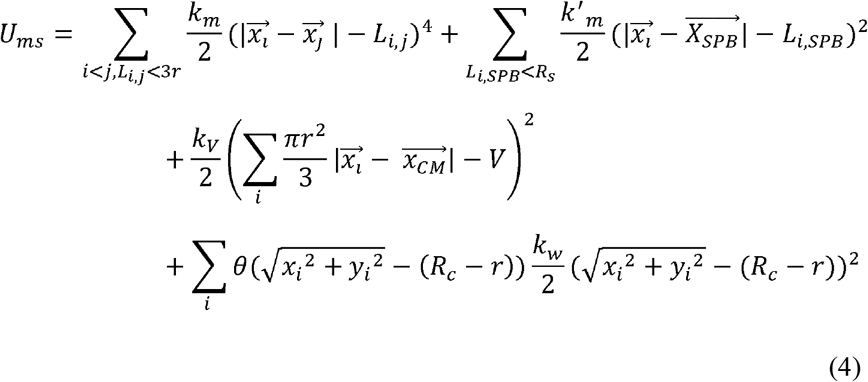

where the *i*-th particle is a part of the nuclear envelope. *θ* is Heaviside step function and *L_ij_* indicates the initial distance between the centers of the *i*-th and *j*-th particles constructing the nuclear envelope (*L_i,SPB_*, the initial distance between the centers of the *i*-th particle and the SPB sphere). Here, the initial structure of the nuclear envelope is indicated as a spherical shell with a thickness of one particle layer constructed by the rings along with *y*-axis with various radii consisting of particles (Fig. 1(c)), and initial position of the center of the SPB sphere is denoted as 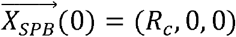 (*R_C_* is the radius of the cell as described below). Σ in the 1st term of *U_ms_* indicates the sum for the *i*-th and *j*-th particles (*i* < *j*) that are adjacent and bonded obeying *L_ij_* < 3*r*; in the 2nd term indicates the sum for the *i*-th particle obeying *L_i,SPB_* < *R_s_*; Σ in other terms indicate sum for all particles constructing the nuclear envelope. Notably, we assumed that the nuclear envelope can deform easily since it consists of lipids that can exhibit the elasticity as well as the fluidity. Thus, we used an unharmonic potential in the 1st term of *U_ms_* which provides weaker tension for local small changes in relative positions among particles in the nuclear envelope than the harmonic potential, inducing larger global deformation of nuclear envelope.

The 2nd term of *U_ms_* indicates the binding of the SPB to the nuclear envelope, where particles constituting the nuclear envelope and confined in the SPB sphere were connected tightly to the center of the SPB sphere. The 3rd term of *U_ms_* provides the force to sustain the volume of the cell nucleus where 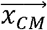 and *V* indicate the position of the center of mass and the volume of the nucleus, respectively. Here, *V* is denoted by 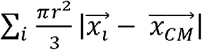 with initial values of 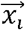. The 4th term of *U_ms_* indicates the excluded volume effect of the cell wall, where the cellular shape was assumed to be a parallel cylinder along the *z*-axis, with a radius *R_C_*. Here, the central axis of this cylinder is denoted as (0,0,*z*), and *θ* indicates the Heaviside function.

*U_e_* denotes the potential of the excluded volume effects among particles and the SPB sphere as follows:

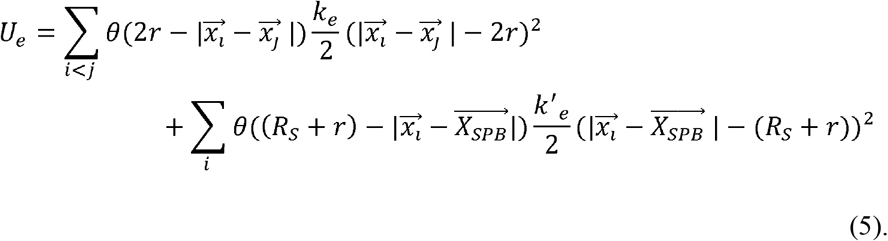

In the third term of equation (1), 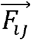 indicates the force of hydrodynamic effects from the *j*-th particle to the *i*-th particle, providing a drag force to each particle from the nucleoplasm. Herein, we employ the lubrication approximation [14, 15] because it can describe hydrodynamic effects among polymers modeled by particle chains in the simulation of the polymers suspension in shear flow suitably [15], which is denoted as follows:

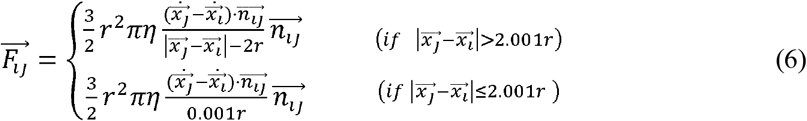

where *r* and 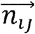denoted the radius of the particle and a relative unit vector from the *j*-th particle to the *i*-th particle 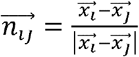, respectively. Nucleoplasm viscosity was assumed the same as the cytoplasm viscosity *η* in this model.

## Parameters for simulations

In the simulations, we assumed a particle radius *r* = 0.084 × 10^−6^(*m*), where each particle models a local region of chromatin of ~100 kb. Here, chromosomes 1, 2, and 3 (5.6 Mb, 4.6 Mb, and 3.5 Mb, respectively) were described by the chains consisting of 56, 46, and 34 particles (*N_1_* = 56, *N_2_* = 46, and *N_3_* = 34; we considered *N_3_* as an even number for convenience of analysis). The nuclear envelope was constructed by using 994 particles (*M*= 994) where we assumed their radii were the same as those of chromosome particles (*r* = 0.084 × 10^−6^(*m*)) because of the convenience to simulate the hydrodynamic effects using Eq. (6). Here, the inner diameter of the initial spherical structure of the nuclear envelope was assumed to be 2.7 × 10^−6^(m), the initial distance between the center of the SPB sphere and that of the nucleus was assumed to be 2.0 × 10^−6^(m), and the radius of the SPB sphere was assumed *R_s_* = 10*r* (Fig. 1(b)). In this case, the diameter of the cross-section where nucleus and the SPB sphere intersect was ~1.0 × 10^−6^(*m*). The initial configurations of the chromosomes were randomized in this spherical shell.

The viscosity of nucleoplasm and cytoplasm *η* was ~0.64 (*kg m*^−1^ *s*^−1^) [12]. The elastic constants were assumed as appropriate large values for the excluded volume effects of all particles and the connections among particles in chromosomes, among particles in chromosomes and the SPB sphere, and among particles in the nuclear envelope and SPB sphere, as *k_c_* = *k_w_* = *k_e_* = *k*′_*e*_ = 2.77 × 10^8^(*kg s*^−2^), *k*′_*c*_ = and. We also assumed the elastic constant to keep the connection of particles in the nuclear envelope and the nuclear volume as appropriate values as *k_m_* = 4.77 × 10^20^(*kg s*^−2^ *m*^−2^) and 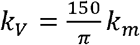 with which the present model reproduced a similar shape variation of the nuclear envelope via horsetail movements. The inertia term of Eq. (1) cannot be eliminated in order to employ the lubrication approximation although the equation of motion of such microscopic system should obey the equation with overdamp limit. However, we assumed *m* as ~*m*/6*πrη* ~ 10^−3^ sec (*m* = 8.3 × 10^−10^(*kg*) was assumed in this paper) that indicates the relaxation times of the particle velocities were negligibly smaller than the period of horsetail movement ~ 10 ^2^ sec. Therefore, we could neglect the effects of inertia to simulate our focused phenomena. We confirmed that the following results were qualitatively unaffected by small changes in these parameters when *k_m_* was chosen appropriate to exhibit sufficient shape variation of the nuclear envelope where chromosomes can elongate. Thus, we consider these parameters appropriate to simulate intracellular environment.

## Results

To evaluate the effects of the horsetail movement, we simulated the following four cases of motions of the SPB sphere in the cylinder (cell) with radius *R_c_* = 2 × 10^−6^(*m*) as shown in Fig. 2: (a) 3-d 8-shaped oscillatory motions as 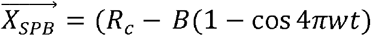, *C*sin4*πwt, A* cos 2*πwt*), (b) 3-d O-shaped oscillatory motions as 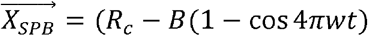, *C*sin 2*πwt, A* cos 2*πwt*), (c) 1-d motions as 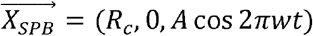, and (d) 1-d constant velocity linear motion as 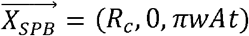. Herein, *A* = 7.3 × 10^−6^(*m*) and 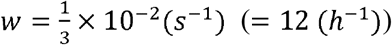 since the SPB completes 30 oscillations in 2 ~3 h from end to end in spheroid-shaped *S. pombe* cells [9]. Furthermore, we assumed *B* = 0.208 × 10^−6^(*m*), and *C* = 0.67 × 10^−6^ (*m*). The results were qualitatively independent of the detail values of *B* and *C*.

**FIG. 2.**
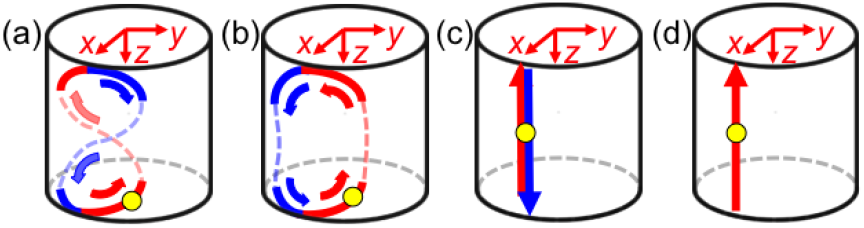
Illustration of trajectories of the spindle pole body (SPB) sphere motions in the cell (cylindrical space). (a) 3-d 8-shaped oscillatory motions, (b) 3-d O-shaped oscillatory motions, (c) 1-d oscillatory motions, and (d) 1-d constant velocity linear motion. Yellow particle indicates the SPB sphere; red and blue arrows indicate the trajectory of downward and upward motion, respectively.

Chromosomal behavior in the migrating nucleus was simulated under the aforementioned 4 conditions. Simulations were repeated with eight initial states for each condition (Figs. 3, 4, and Supplementary movies 1 and 2). Figure 3 shows typical snapshots for t = 75, 3375, and 7950 (s) in an example of 8-shaped 3-d oscillatory motion. Elongation of the nuclear envelope (Supplementary movie 1) and switching of relative positions among chromosomes (Fig. 3, Supplementary movie 2) were observed during oscillatory motions of the SPB. The present results indicate the validity of our simulations.

**FIG. 3.**
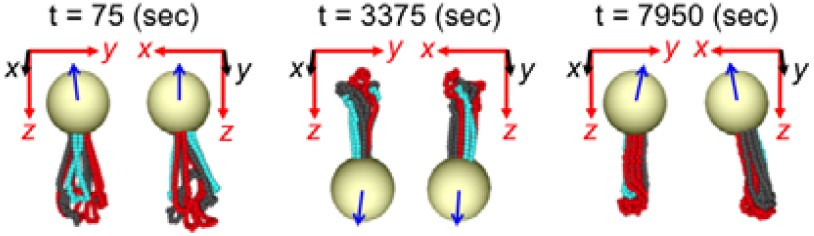
Typical time course of chromosome dynamics in the case that the spindle pole body (SPB) sphere exhibits 3-d 8-shaped oscillation. Snapshots of chromosome configurations in the nucleus from two points of view (projected on *y-z* and *x-z* plane) at time t = 75 s (after 0 oscillations), 3375 s (after 10 oscillations), and 7950 s (after 26 oscillations). The large sphere indicates the SPB sphere; red, black, and blue particle chains indicate Chr. 1, Chr. 2, and Chr. 3, respectively. Blue arrows indicate the moving directions of SPB.

To obtain a pictorial view of chromosomal movements, we plotted positions of the chromosomes as probability distributions along the x-y plane (Fig. 4). The plots were obtained from the 34 (= *N_3_*) nearest particles from the SPB sphere (*N_3_* / 2 = 17 nearest particles from each end) along each of the 6 chromosomes (Fig. 4a). Figure 4b shows typical snapshots of the probability distributions of particles for t = 75, 3375, and 7950 (s) in the 3-d 8-shaped oscillatory motion. When t = 75, the probability distributions of particles of chromosome 1 and 3 exhibited two separated hills respectively, that indicated homologous pairs of chromosomes 1 and 3 were separated far away. On the other hand, the probability distributions of particles of all chromosomes exhibited only one hill respectively at different area when t = 7950. These facts indicated each homologous chromosome pair gathered temporally and non-homologous chromosomes were separated from each other, that is, alignment of homologous chromosomes occurred.

**FIG. 4.**
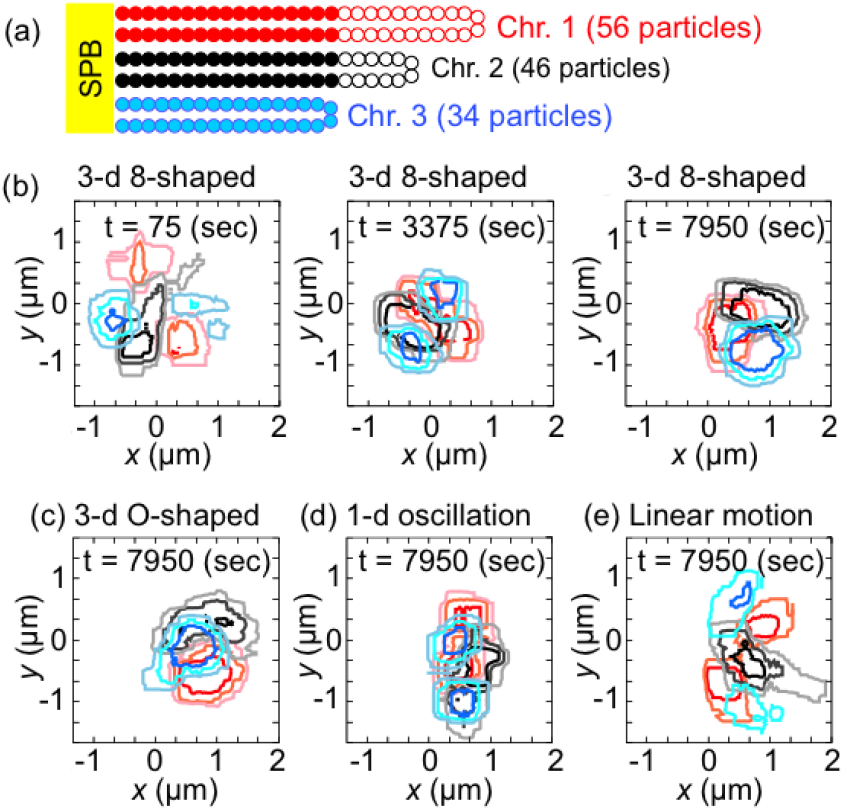
Probability distributions of particles in each of the chromosome chains near the spindle pole body (SPB) sphere along the *x-y* plane. (a) The 34 (= *N_3_*) nearest particles from the SPB sphere along each of the 6 chromosomes (filled particles) were used to estimate probability distributions. (b) Probability distributions of the 34 (= *N_3_*) nearest particles on each chromosome indicated as filled particles in (a). Curves represent contour lines of probability distributions of particles in the 3-d 8-shaped oscillation of the SPB sphere at t = 75, 3375, and 7950 s; red, black, and blue curves indicate Chr. 1, Chr. 2, and Chr. 3, respectively. The intensity of the contour lines represents the probability values of 0.3, 0.5, and 0.75 in an increasing order of darkness. (c) – (e) Probability distributions at t = 7950 s in (c) 3-d O-shaped oscillation, (d) 1-d oscillations, and (e) constant velocity linear motion of the SPB sphere. Colors of the contour lines represent probability distributions of each chain as in (b).

Figure 4 (c) – (e) show typical snapshots of the probability distributions of particles at t = 7950 (s) when the SPB sphere exhibits a 3-d O-shaped oscillatory motion, a 1-d oscillatory motion, and a linear motion. As shown in Fig. 4 (b) and (c), each homologous pair of chromosomes gathered temporally and non-homologous chromosomes were separated from each other when the SPB sphere exhibits 3-d oscillatory motion involving 3D reversal (torsional turning) motion of chromosomes.

However, when the SPB sphere exhibits 1-d oscillatory motion involving no torsional motion of chromosomes, some homologous chromosome pairs often stay separated from each other (Fig. 4 (d); alignment of homologous chromosomes was observed exceptionally only in one out of eight simulations). Furthermore, the assembly of homologous chromosomes did not occur when the SPB sphere exhibits a linear motion involving no chromosomal turning (Fig. 4 (e)). These results suggest that chromosomal torsional turning motion is crucial for shuffling of chromosomes for appropriate alignment of homologous chromosomes.

By the simulations of following virtual situations where chromosomes 1 and 2 became shorter as (i) 4.6 Mb and 4.0 Mb (*N_1_* = 46, *N_2_* = 40, and *N_3_* = 34), and (ii) 3.8 Mb and 3.6 Mb (*N_1_* = 38, *N_2_* = 36, and *N_3_* = 34), we found that the frequency of the separation of non-homologous chromosomes decreased with the decrease in their difference in chain lengths: the separation of non-homologous chromosomes was observed in three out of eight simulations in both situations even when the SPB sphere exhibits a 3-d 8-shaped oscillatory motion. We also found the separation of non-homologous chromosomes occurred little if the elastic constant *k_m_* among particles in nuclear envelope was given so large as that the shape variation of the nuclear envelope was not sufficient for the elongation of chromosomes via the SPB movements. By these facts, non-homologous chromosomes are considered to separate owing to differences in chain lengths among non-homologous chromosomes.

During the horsetail movement, the SPB movement reverses its direction periodically. Chromosomes receive drag forces from the nucleoplasm at every turn of the SPB, while no drag force is exerted on chromosomes and the nuclear envelope if SPB migrates linearly with a constant velocity. During reversal of SPB, the region distal from the SPB in longer chromosomes tends to be dragged by the slow (nearly stopped) motion of the rear region of the bent nucleus, while that in the shorter one tends to follow the SPB motion (Fig. 5). Thus, the relative positions of longer and shorter chromosomes tend to separate gradually from each other owing to reversal of the iterative direction of the SPB movement, resulting in the separation of non-homologous chromosomes. Owing to this separation, homologous chromosomes are aligned and stabilized during the periodic reciprocation of the SPB.

**FIG. 5.**
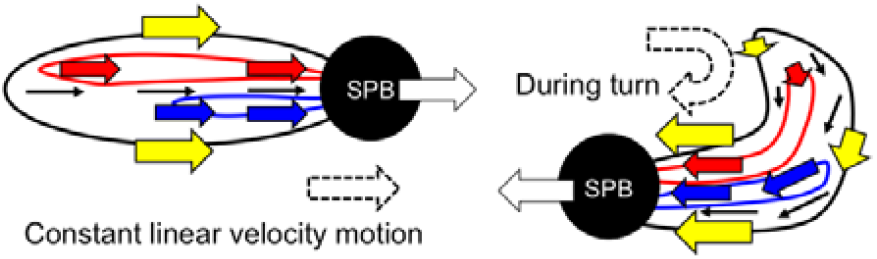
Illustration of velocities of the spindle pole body (SPB; white arrow), long and short chromosomes (red and blue arrows, respectively), the nuclear envelope (yellow arrows), and the nucleoplasmic flow (black arrows). When the SPB exhibits a constant linear velocity motion, all regions of the nuclear envelope, the nucleoplasm, and chromosomes relax to the same velocity as the SPB (Left). During the turn of the SPB, the regions of the nuclear envelope, the nucleoplasm, and chromosomes at the nuclear front end (near the SPB) follow the motion of the SPB, while those at the nuclear rear end are almost stopped (Right). In the latter case, longer chromosomes tend to be dragged by the slow motion of the nuclear rear region, while shorter chromosomes tend to follow the motion of the SPB.

## Discussion

Homologous chromosome pairing has received increasing attention in studies in physics, and several models have been proposed to describe its underlying mechanisms [10, 11, 12, 13, 16]. In the present simulations, separation of non-homologous chromosomes was achieved when the SPB sphere shows 3-d oscillations in the cell, but not when the SPB sphere exhibits only a linear motion. This result is further substantiated by previous experimental evidence regarding *S. pombe:* the efficiency of homologous recombination decreases drastically when the nucleus does not exhibit the horsetail movement but just elongates [17]; the iterative reversals of the direction of the SPB movement during the horsetail movement are essential for the alignment of homologous chromosomes via separating non-homologous chromosomes [7, 9]. Our simulation reveals that a major force for the alignment and sorting of homologous/non-homologous chromosomes is the nucleoplasmic flow generated owing to SPB reversal and deformation of the nuclear envelope, involving direct collision and indirect hydrodynamic interactions between chromosomes and the nuclear envelope, which were previously unclear. The present model is the first, to our knowledge, to consider both direct collisions and indirect hydrodynamic interactions as essential mechanisms for the alignment of homologous chromosomes.

This study was partly supported by the Platform Project for Support in Japan Agency for Medical Research and Development (to AA); MEXT KAKENHI Grant Number JP 17K05614 (to AA), JP18H05528 (to TH) and JP18H05533 (to YH).

